# Radiation-induced interferon-I response impairs thyroid organoid function

**DOI:** 10.64898/2026.01.14.699489

**Authors:** Rufina Maturi, Davide Cinat, Anne L Jellema-de Bruin, Gabriella De Vita, Schelto Kruijff, Rob P. Coppes, Abel Soto-Gamez

**Affiliations:** Department of Molecular Medicine and Medical Biotechnology, University of Naples Federico II, Naples, Italy; Department of Biomedical Sciences, University Medical Center Groningen, University of Groningen, Groningen 9713 GZ, The Netherlands; Department of Radiation Oncology, University Medical Center Groningen, University of Groningen, Groningen 9713 GZ, The Netherlands; Department of Surgery, University Medical Center Groningen, University of Groningen, Groningen 9713 GZ, the Netherlands; Department of Nuclear Medicine and Molecular Imaging, University of Groningen. University Medical Center Groningen, Groningen, The Netherlands; Department of Molecular Medicine and Surgery, Karolinska Institutet, Stockholm, Sweden

## Abstract

**Background and Aim:** Radiotherapy is a standard cancer treatment, but radiation exposure to surrounding healthy tissues may lead to adverse side effects that compromise patient quality of life. In patients with head and neck cancer treated with radiotherapy, thyroid damage is a frequent complication, resulting in hypothyroidism or secondary thyroid malignancies. Although clinically recognized, the molecular mechanisms underlying these side effects remain mostly unexplored. This study aims to characterize the radiation-induced molecular alterations in thyroid organoids.

**Methods:** Bulk RNA sequencing was performed to investigate transcriptomic changes in tissue-derived thyroid organoids following gamma-irradiation. Observed changes were further validated and explored using qPCRs, western blotting, immunofluorescence, caspase 3/7 activity and organoid forming efficiency.

**Results:** Our findings identify interferon-β (IFN-β) signaling as a key mediator of radiation-induced inflammation in the thyroid. Additionally, the intrinsic apoptotic pathway was found to be the predominant mechanism of radiation-induced thyroid cell death. Notably, while IFN-β exhibited a protective effect against apoptosis, it concurrently reduced thyroid stem progenitor cell potential.

**Conclusions:** These results highlight the dual role of IFN-β signaling in modulating thyroid cell fate after irradiation, potentially promoting survival upon injury at the expense of regenerative potential.

## Introduction

Head and neck cancer (HNC) is the seventh most common malignancy worldwide [1]. The first-line treatment for most patients includes radiotherapy, often in conjunction with surgery and chemotherapy [1]. Radiation-induced damage to the healthy tissue surrounding the tumor often leads to side effects impairing the patient’s quality of life [2]. Hypothyroidism is one of the most common side effects associated with curative radiotherapy for HNC [3]. The thyroid gland is the largest purely endocrine gland in humans, and it secretes two main thyroid hormones (triiodothyronine; T3 and thyroxine; T4), which are crucial in normal growth, development, and metabolism regulation, affecting all organ systems [4]. Within five years after radiation exposure, 36% of patients develop hypothyroidism and are left with the lifelong need for substitutive therapy [3]. The first reports on hypothyroidism following radiotherapy for HNC were published in the 1960s; nevertheless, the molecular mechanisms underlying thyroid damage remain largely unknown [5–6]. Moreover, therapeutic and incidental radiation exposure increases the risk of thyroid tumors. These are approximately 70% benign lesions, while the remaining 30% are malignant, mostly well-differentiated thyroid carcinomas with few fatal cases [7].

Radiation exposure causes DNA damage, which, next to cell death, activates a cascade of events that, among others, can trigger inflammation [8]. While the physicochemical events subside quickly after radiation exposure (e.g., free radical generation), the inflammatory response persists, resulting in the release of cytokines, chemokines, and growth factors that alter the tissue physiology [9] and regenerative capability of the surviving cells [8]. Most studies exploring the events that occur after radiotherapy have focused on tumor tissues [10]; nevertheless, the activation of similar responses in healthy tissues may underlie side effects and warrants further investigation.

Interferon-beta (IFN-β), a type-I interferon (IFN-I), exhibits immunomodulatory and anti-proliferative properties that may enhance the therapeutic efficacy of radiotherapy in cancer treatment [11]. Preclinical studies suggest that IFN-β can sensitize tumor cells to ionizing radiation by modulating DNA damage response pathways and altering the tumor microenvironment [12]. Additionally, IFN-β may potentiate immune-mediated tumor clearance through upregulation of MHC class I expression and the activation of cytotoxic T cells. The synergistic interaction between IFN-β and radiotherapy holds promise for improving tumor control and reducing recurrence. However, clinical reports highlight the toxic effects of IFN-I treatments on the thyroid, mainly as an association with autoimmune thyroid diseases, which can also lead to hypothyroidism [13–16].

Organoids are three-dimensional (3D) structures that can be grown starting from tissue-isolated cells, as in this study, or *via* the differentiation of pluripotent stem cells (iPSCs or ESCs) [17–22]. As such, organoids are useful models to study acute and early radiation effects in specific tissues. In this study, we used mouse thyroid gland organoids (mTGOs) resembling the tissue of origin [17] to characterize the molecular events occurring in thyroid gland cells following radiation exposure. Herein, we show that irradiation is associated with a pronounced inflammatory response as indicated by the enhancement of IFN-β signaling.

## Materials and Methods

### Isolation of Murine Thyroid Cells and Organoid Culture

All animal procedures were approved by the Ethical Committee for Animal Testing at the University of Groningen. Thyroid glands were harvested from female C57BL/6 mice aged 8–12 weeks (Harlan, The Netherlands). Tissues were collected in Hank’s Balanced Salt Solution (HBSS; Gibco) supplemented with 1% bovine serum albumin (BSA) and subjected to mechanical dissociation using the GentleMACS Dissociator (Miltenyi Biotec). This was followed by enzymatic digestion in HBSS/1% BSA containing calcium chloride (6.25 mM), collagenase I (100 U/mL; Gibco), and dispase (1.5 U/mL; Gibco). The resulting cell suspension was cultured overnight in DMEM/F12 medium (Gibco; #11320-074) supplemented with 1% penicillin/streptomycin (Gibco; #15140-163), 2 mM GlutaMAX (Gibco; #35050038), epidermal growth factor (EGF; 20 ng/mL; Sigma-Aldrich; #E9644), fibroblast growth factor 2 (FGF-2; 20 ng/mL; Peprotech; #100-18-B), and 0.5% B27 supplement (Gibco; #17504044). After 24 hours, cell aggregates were collected, enzymatically dissociated into single cells using 0.05% trypsin-EDTA (Gibco), and filtered. Cells were resuspended in DMEM/F12, counted, and seeded at a density of 10,000 cells in a 25 µL volume, which was mixed on ice with 50 µL of Basement Membrane Matrigel^TM^ (BD Biosciences; #356235). The cell-Matrigel mixture was plated in the center of 12-well culture plates. After Matrigel polymerization (30 minutes at 37°C), 1 mL of thyroid growth medium (TGM) was added. TGM consisted of DMEM/F12 supplemented with 1% penicillin/streptomycin, 2 mM GlutaMAX, EGF (20 ng/mL), FGF-2 (20 ng/mL), ROCK inhibitor Y-27632 (10 µM; Abcam; ab120129), and 0.5% B27 supplement.

### Organoid Self-Renewal Assay

Organoid-forming capacity was assessed 7– or 11-days post-seeding (2 or 4 days after irradiation). Secondary spheres were counted, and the sphere-forming efficiency was calculated. For passaging and self-renewal capacity, organoids were released from the Matrigel using dispase (1 mg/mL), dissociated into single cells with 0.05% trypsin-EDTA, and reseeded in Matrigel^TM^ as described above, using 10,000 per well. Organoid counts and total cell numbers were recorded to determine organoid formation efficiency (OFE) and population doublings (PD) using the following formulas:

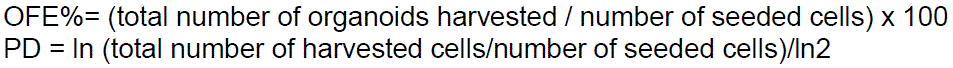

### Organoid irradiation and treatments

Either single cells seeded after isolation or 5-day-old thyroid organoids were irradiated with 2,4, 7 and 15 Gy of photon irradiation, using a Cesium-137 source (IBL 637) with a dose rate of 0.59 Gy/min at the department of Biomedical Sciences of the UMCG. IFN-β (MedChemExpress, #HY-P73130) was added to the media of gamma-irradiated and non-irradiated organoids at day 7 at a final concentration of 2 ng/mL and at the time of self-renewal.

### Mice irradiation and treatments

Thyroid glands of 12-week-old female C57BL/6 mice (Envigo, the Netherlands) were (sham-) irradiated with a single dose of 15 Gy X-rays (RAD 320, Precision X-ray) and sacrificed 6 days after irradiation.

### RNA Isolation and Quantitative PCR Analysis

Total RNA was extracted from thyroid organoids using the RNeasy Mini Kit (Qiagen; #74104), according to the manufacturer’s protocol. Complementary DNA (cDNA) was synthesized from isolated RNA using a reverse transcription reaction comprising 1 µL of dNTPs (10 mM; #10297–018), 1 µL of random primers (100 ng; #SO142), 4 µL of 5× First-Strand Buffer (#28025013), 2 µL of dithiothreitol (DTT, 0.1 M; #328025013), 1 µL of RNaseOUT™ recombinant ribonuclease inhibitor (40 U/µL; #10777019), and 1 µL of M-MLV reverse transcriptase (200 U; #28025013). Reagents were purchased from Invitrogen. Quantitative PCR (qPCR) was performed using gene-specific primers and iQ SYBR Green Supermix (Bio-Rad, #170–8885) on a Bio-Rad Real-Time PCR Detection System. All reactions were conducted in technical triplicates. Gene expression levels were normalized to *Ywhaz*, which served as the internal reference gene. The sequences of all primers used are listed in Supplementary Table 1.

### Bulk RNA Sequencing and Analysis

Total RNA was isolated from thyroid organoids cultured for 7, and 11 days, with and without irradiation treatment (see as described above). RNA integrity was assessed using High Sensitivity RNA ScreenTape assays on the Agilent 2100 Bioanalyzer, and only samples with a RNA integrity number (RIN) greater than 9 were used for downstream applications. RNA samples from organoids derived from three individual animals were subjected to cDNA library preparation using the Smart3SEQ protocol [35] following the manufacturer’s instructions. For each sample, 80 ng of total RNA was used as input. Resulting libraries were pooled at equimolar concentrations and sequenced on an Illumina NextSeq 500 platform at the ERIBA Research Sequencing Facility (University Medical Center Groningen, UMCG). A 75 bp single-end sequencing run was performed, using a final pool concentration of 1.8 pM with 15% PhiX control library spike-in. Sequencing reads were aligned to the *Mus musculus* reference genome (GRCm39) using the STAR aligner. Gene-level expression values were normalized, and Z-scores for selected genes of interest were manually calculated.

### Protein Extraction and Western Blot Analysis

Organoids were released from the Matrigel matrix using dispase (1 mg/mL), then lysed in radioimmunoprecipitation assay (RIPA) buffer. Lysates were sonicated and centrifuged at 4°C for 5 minutes at maximum speed to remove debris. Protein concentration was determined using the Bradford assay (Bio-Rad), and equal amounts of total protein were denatured by boiling at 99°C for 5 minutes in sample buffer.

Proteins were separated on 12% SDS-polyacrylamide gels and transferred onto nitrocellulose membranes using the Trans-Blot Turbo Transfer System (Bio-Rad). Membranes were blocked in 10% non-fat milk in PBS-Tween 20 (PBS-T) for 1 hour at room temperature, followed by overnight incubation at 4°C with primary antibodies diluted in 3% BSA in PBS-T. After washing, membranes were incubated with horseradish peroxidase (HRP)-conjugated secondary antibodies for 1.5 hours at room temperature. Signal detection was performed using enhanced chemiluminescence (ECL) reagents, and membranes were imaged using the ChemiDoc Imaging System (Bio-Rad). Band intensities were quantified using Image Lab software (Bio-Rad). The following primary antibodies were used: GAPDH (1:10,000; Fitzgerald, #10R-G109a), STAT1 (1:1000; Cell Signaling, #14994T), RIG-I (1:600; Santa Cruz Biotechnology, #sc-376845), phospho-STAT1 (Tyr701) (1:600; Cell Signaling, #9167S). HRP-conjugated secondary antibodies (GE Healthcare, NA934, #NXA931V) were used at 1:3,000 dilution, all prepared in 3% BSA in PBS-T.

### Incucyte-Based Thyroid *Organoid Size Analysis*

Mouse thyroid organoid cells were seeded into 24-well-plates and cultured in Matrigel^TM^ domes in TGM media, inside an IncuCyte® S3 live-cell imaging system (Essen BioScience) as previously described [26]. Plates were kept at 37 °C in a humidified atmosphere with 5% CO₂. Imaging was performed using the organoid module, capturing phase contrast + brightfield, with a 4x objective (S3/SX1 G/R Optical Module). The subsequent analyses were made using a 200 µm radius, 50 units of sensitivity, edge split sensitivity of 50 units, and a hole fill cleanup size of 5E+0-05 µm2. Measurements were performed on days 0, 4, 7 and 10. For comparisons between conditions, relative organoid size was calculated by normalizing total object area from each sample by its own measurements on day 0.

### Caspase 3/7 assays

Mouse thyroid organoids were harvested from 24-well-plates at Day 5 and irradiated at various doses, before being transferred into (untreated) 96 well plates in DMEM:F12 media with a fluorogenic substrate (1 µM Cell Event Caspase 3/7 Green, ThermoFisher Scientific). Caspase 3/7 activity was measured based on fluorescence intensity recordings performed every 2 hours in an IncuCyte® S3 live-imaging system using whole well scans under 10X magnification and 300ms exposure in the green channel. The Total Green Object Fluorescent Intensity was calculated for each well at every timepoint. The area under the curve (AUC) was calculated for each condition for up to 48 hours, using GraphPad Prism 8.0.2, and a baseline centered at 0. Outlier exclusion criteriz were based on the mean fluorescent intensity across technical replicates, using an upper and lower fluorescent threshold of μ ± 2.0 σ.

### Organoid fixation and immunofluorescence

Thyroid gland organoids were harvested and fixed in 4% formaldehyde for 15 minutes, followed by embedding in HistoGel. The embedded samples were dehydrated, paraffin-embedded, and sectioned into 4 µm slices. Sections were dewaxed and subjected to heat-induced antigen retrieval using Tris-EDTA buffer (pH 9). Permeabilization and blocking were carried out using a blocking solution comprising 4% goat serum, 1% BSA, and 0.01% Triton X-100 in 1× PBS. Sections were then incubated overnight at 4 °C with the following primary antibodies: STAT1 (Cell Signaling, #14994T, 1:100), MX1 (Proteintech, #13750-1-AP, 1:100). The next day, secondary antibody incubation with Alexa Flour 488 goat anti-rabbit (ThermoFisher Scientific, #A-11008, 1:1000) was performed for 1 hour at room temperature, followed by nuclear staining with DAPI for 10 minutes. Slides were mounted and imaged using a Leica DM6 microscope. Image analysis was conducted using ImageJ (v1.52).

### Annexin V/Propidium Iodide Staining and Flow Cytometry Analysis

Apoptosis was assessed using the Annexin V-FITC/Propidium Iodide (PI) (Invitrogen, #V13242) double staining method followed by flow cytometric analysis. mTGOs were harvested at D11 after treatment with IFN-β from D5, trypsinised and washed twice with cold phosphate-buffered saline (PBS). Approximately 1 × 10⁶ cells were resuspended in 100 µL of 1X Annexin V binding buffer. Annexin V-FITC (5 µL) and PI (5 µL of a 50 µg/mL stock solution) were added to each sample, gently mixed, and incubated for 15 minutes at room temperature in the dark. After incubation, 400 µL of binding buffer was added to each tube, and samples were immediately analyzed by flow cytometry using Quanteon flow cytometer at the Flow Cytometry Unit of the UMCG. A total of 100,000 events per sample were collected. Data were analyzed using FlowJo™ software (v10.7.2; Tree Star, Inc.). Compensation controls were performed using single-stained Annexin V-FITC and PI samples. Cells were gated to exclude debris (FSC-A, SSC-A) and doublets (FSC-A, FSC-H), using the same gates for all samples. Four distinct populations were identified: live cells (Annexin V⁻/PI⁻), early apoptotic (Annexin V⁺/PI⁻), late apoptotic or necrotic (Annexin V⁺/PI⁺), and necrotic cells (Annexin V⁻/PI⁺).

### Statistical Analysis

Statistical analyses were performed using GraphPad Prism (version 8). Differences between groups were assessed using two-tailed unpaired Student’s *t*-tests or one-way ANOVA followed by Tukey’s post hoc test for multiple comparisons, as appropriate. Data are reported as mean ± standard error of the mean (s.e.m.). Biological replicates (*N*) represent independent samples derived from distinct animals, with the specific number of replicates and statistical tests detailed in the corresponding figure legends. Statistical significance was defined as *p* ≤ 0.05.

## Results

The molecular basis for thyroid gland radiosensitivity has not been explored, despite apparent clinical deleterious effects [3], [6], [22]. Therefore, we used primary thyroid cells isolated from murine thyroid glands directly after enzymatic digestion as a starting point to gain better insight into the cellular events occurring in the thyroid after radiation exposure. Hereto, we gamma-irradiated dispersed murine thyroid gland cells to different doses (0, 2, 4, 7, and 15 Gy). Next, (sham)-irradiated primary thyrocytes were seeded in Matrigel^TM^ domes to form organoids (Fig. 1A). As organoids are complex multicellular structures that recapitulate the tissue of origin, their development is the result of an intricate combination of proliferation and differentiation [23]. Therefore, the ability of a single thyroid cell to form organoids can be defined as an indirect measurement of its stemness [17]. As such, we quantified the number of organoids formed by primary thyroid gland cells, one week after being seeded in Matrigel^TM^ domes directly after irradiation, hereafter referred to as Organoid Forming Efficiency (OFE) (Fig. 1A). Irradiation resulted in a dose-dependent decrease in OFE (Fig. 1B, C), which was completely abolished after 15 Gy (Fig. 1C). This data highlights thyroid gland radiosensitivity compared to other glandular organs, such as the salivary gland. and is further supported by the observation that cells isolated from the thyroids of 14 Gy-irradiated mice were unable to grow any complex structures *ex vivo* (Supp. Figure 1A and B). Therefore, this dose was excluded from further analyses, as it completely sterilized tissue-specific stem-like cells and did not yield enough material to explore the molecular processes involved.

**Figure 1.**
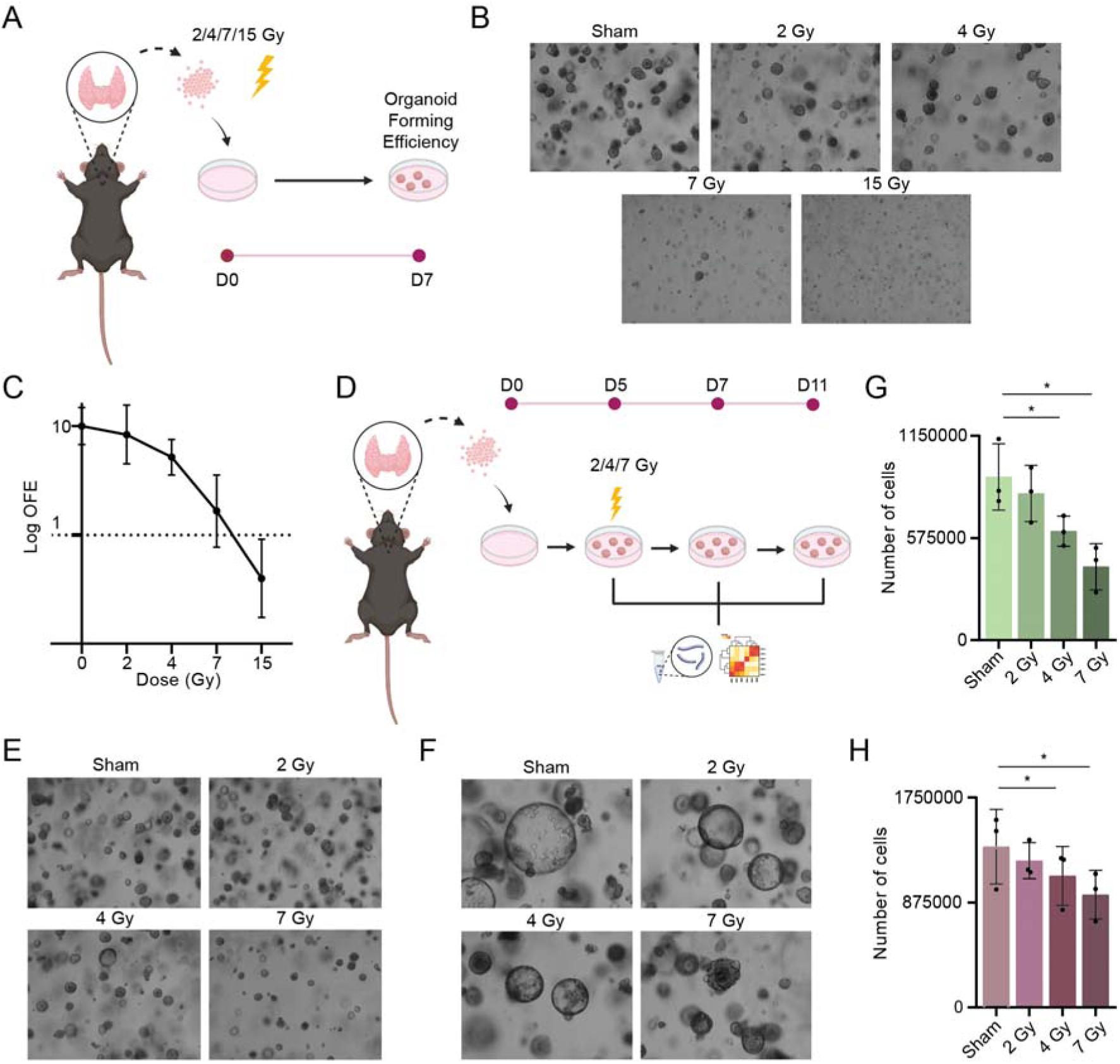
Radiation exposure impairs primary thyrocyte stemness and organoid formation. (A) Experimental setup for isolated thyroid cell irradiation: cells are isolated from mice thyroid glands by enzymatic and mechanical digestion, seeded in a solid Matrigel^TM^ dome, irradiated with 0 (sham), 2, 4, 7 or 15 Gy, and collected after 7 days. (B) Light microscopy images of organoids grown from irradiated cells for 7 days, 4x objective. (C) Dose response curve of the relative OFE calculated 7 days after irradiation of single cells with 2,4,7 and 15 Gy. (D) Experimental setup for irradiation of mTGOs: cells were isolated from mice thyroid glands by enzymatic and mechanical digestion and seeded in Matrigel^TM^ domes. 5-day-old organoids were irradiated with 0 (sham), 2, 4 and 7 Gy and harvested at D7 and D11. (E) Light microscopy images of D7 organoids irradiated at D5 with 0, 2, 4 and 7 Gy, 4x objective. (F) Light microscopy images of D11 organoids irradiated at D5 with 0, 2, 4 and 7 Gy, 4x objective. (G) Total number of cells counted after trypsinization of organoids at D7. (H) Total number of cells counted after trypsinization of irradiated organoids at D11. Data are shown as mean ± SEM. In all panels, group differences were evaluated using one-way ANOVA with post-hoc Tukey’s HSD. n = 3 animals/condition. \**p* < 0.05

Despite providing important information about thyrocyte radiosensitivity, the irradiation of single cells fails to mimic the complex, multicellular environment of native tissue. Therefore, to better recapitulate alterations occurring *in vivo*, we seeded primary thyrocytes in Matrigel^TM^ domes and allowed them to form organoids. By day 5 in culture (D5), the structures were big enough to be considered as organoids, at which point they were irradiated to 0, 2, 4, or 7 Gy. To assess both acute and early events following radiation, organoids were analyzed at 2 and 6 days after exposure, the equivalent of 7 and 11 days in culture, respectively (Fig. 1D). Representative light microscopy images of day 7 (D7) and day 11 (D11) organoids are shown (Fig. 1E-F). Expectedly, increasing doses of radiation led to a progressive reduction in the total number of cells composing the organoids, both at D7 (Fig. 1G) and D11 (Fig. 1H), suggesting either less proliferation or increased cell death after the treatment.

To investigate the signaling pathways paralleling the response observed after irradiation (Fig. 1C, 1F, 1H), we performed bulk RNA sequencing (RNA-seq) on organoids collected at 2 days (D7) and 6 days (D11) following irradiation. Differential expression analysis revealed clear dose-dependent transcriptional changes at both timepoints (Fig. 2A). Organoids exposed to 7 Gy showed the highest number of DEGs compared to sham-irradiated controls (Fig. 2A), consistent with the pronounced cytotoxic effects observed in Fig. 1B and 1E. For this reason, downstream analyses were performed using 7 Gy only, coupled with previous data demonstrating that 7 Gy induces substantial phenotypic changes that closely recapitulate responses in other glandular organoid models [24–26].

**Figure 2.**
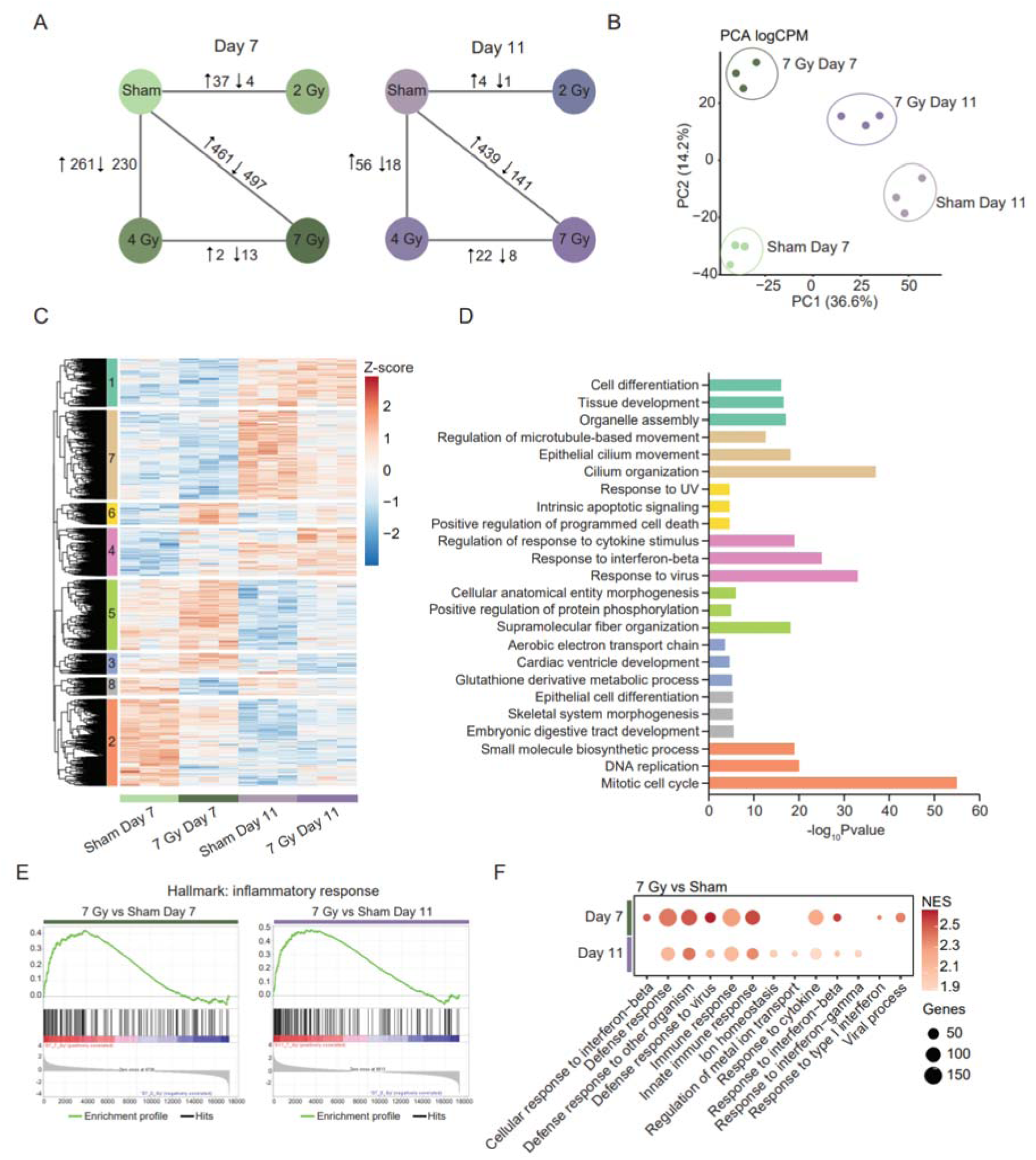
Radiation-induced transcriptional changes. (A) Bulk RNA-seq analysis of Day 7 and Day 11 mTGOs showing the number of significantly up– and downregulated genes across different radiation doses. (B) PCA plot of sham and 7 Gy-irradiated samples at Day 7 and Day 11. Each dot represents a different biological replicate (n = 3 animals per condition). (C) Heatmap of sham and 7 Gy-irradiated samples at Day 7 and Day 11, showing hierarchical clustering. (D) Bar plot showing the top 3 significant biological processes for each cluster identified in (C). Colors correspond to cluster groups in (C). (E) GSEA plots of the hallmark inflammatory response pathway comparing 7 Gy vs. sham samples at Day 7 and Day 11. (F) Dot plot showing the top 10 significant Gene Ontology (GO) terms in Day 7 and Day 11 irradiated organoids compared to sham. In all panels n = 3 animals/condition.

Principal component analysis (PCA) revealed distinct clustering based on both dose and time points (Fig. 2B), indicating substantial transcriptional differences between the conditions. Hierarchical clustering further supported this observation (Fig. 2C), with Day 7 sham organoids displaying an enrichment of cell cycle and stem cell-associated biological processes (Fig. 2D). In contrast, irradiated organoids exhibited a strong upregulation of inflammatory, interferon-related, and apoptotic gene ontology terms (Fig. 2D). Importantly, the impact of interferon-related genes on thyroid function, and whether IFN-I contributes to the observed reduction in stem cell potential, remains uncertain. Indeed, gene set enrichment analysis (GSEA) confirmed a significant induction of inflammation-related genes (Fig. 2E), as well as interferon-β and immune signaling pathways (Fig. 2F) at both Day 7 and Day 11.

To confirm the upregulation of IFN-I signaling after irradiation, we assessed the expression of interferon-stimulated genes (ISGs) and transcription factors downstream of IFN-I release and signaling activation. Bulk RNA-seq and qPCR analyses showed increased expression of several ISGs at both D7 and D11 (Fig. 3A, B). Additionally, immunofluorescence staining and western blot analysis revealed elevated levels of STAT1 and p-STAT1 at D11, whereas p-STAT1 rises already at D7 (Fig. 3C, D, E), which are transcription factors essential for mediating the downstream IFN-responses [27]. Moreover, D7 and D11-irradiated mTGOs showed significantly higher levels of the ISGs MX1 and RIG-I (Fig. 3F, G, H), aligning with previous studies [28] and confirming the upregulation of the IFN-I/STAT1/ISG signaling axis. Overall, these findings confirm strong transcriptional changes after irradiation and suggest IFN-β as a potential mechanism involved in the functional decline of the thyroid gland following irradiation.

**Figure 3.**
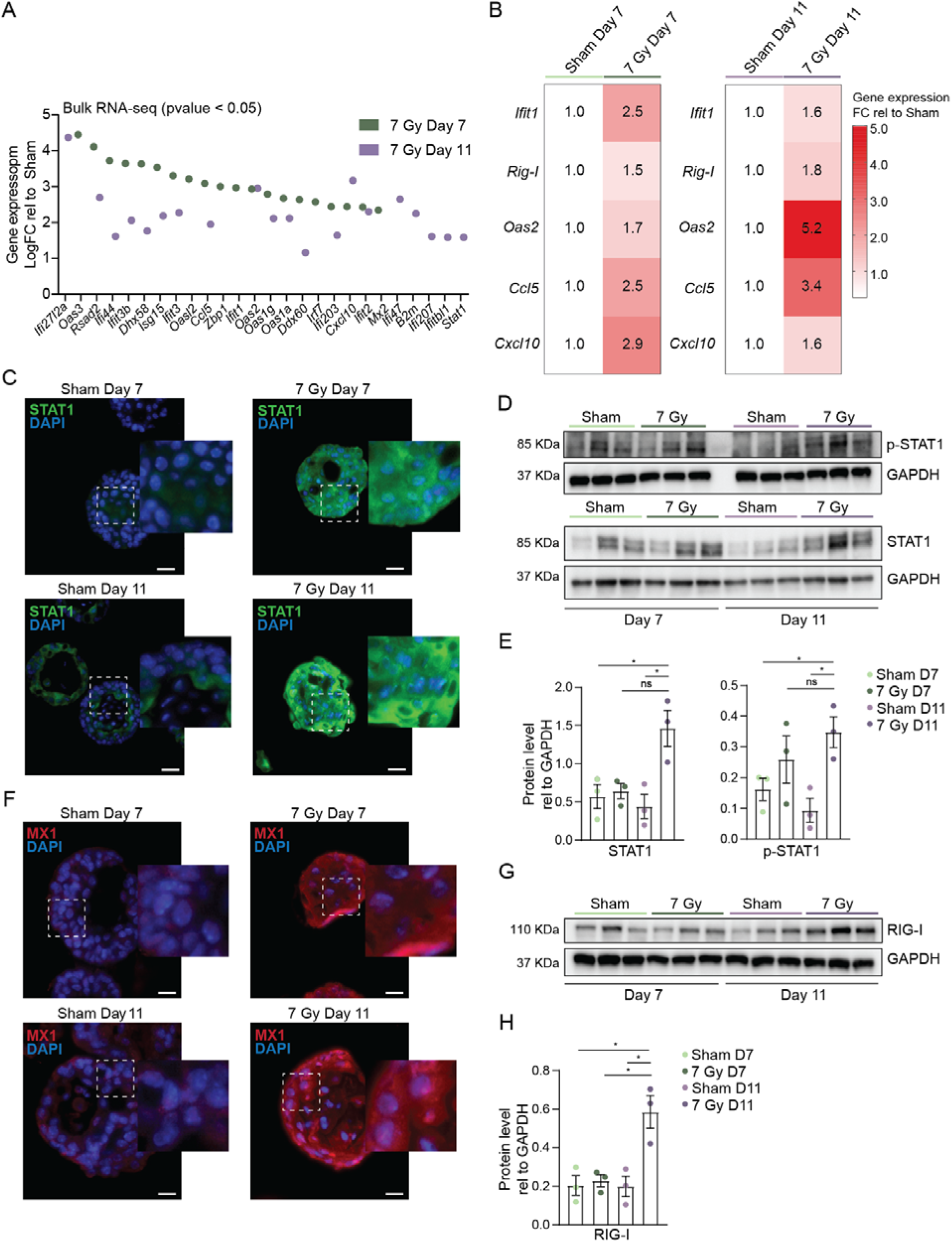
Radiation induces IFN-I signaling upregulation. (A) Top 20 significant ISGs extrapolated from bulk RNA-seq analysis of 7 Gy-irradiated mTGOs at Day 7 and Day 11. (B) rt-qPCR analysis of *Ifit1, Rig-I, Oas2, Ccl5, Cxcl10* of 7 Gy-irradiated mTGOs at Day 7 and Day 11. Data is shown as FC relative to Sham. (C) Representative images of immunofluorescence staining of sham– and 7 Gy-irradiated mTGOs sections at Day 7 and Day 11. (D) Western blot analysis of p-STAT1 and STAT1 of sham– and 7 Gy-irradiated mTGOs at Day 7 and Day 11. GAPDH was used as a loading control. (E) Quantification of the western blot shown in Figure 3D. (F) Representative images of immunofluorescence staining of sham– and 7 Gy-irradiated mTGOs sections at Day 7 and Day 11. (G) Western blot analysis of RIG-I of sham– and 7 Gy-irradiated mTGOs at Day 7 and Day 11. GAPDH was used as a loading control. (H) Quantification of the western blot shown in Figure 3G. In all panels, data are shown as mean ± SEM. Group differences were evaluated using one-way ANOVA with post-hoc Tukey’s HSD. n = 3 animals/condition. *p < 0.05. Scale bar 25 μm.

To investigate the role of IFN-β activated by radiation exposure, a preliminary analysis was conducted to assess the effect of IFN-β on thyroid cell viability. Our RNA-seq data indicated radiation-induced thyroid cell death predominantly occurs via the intrinsic apoptotic pathway and is rescued by pan-caspase inhibition (Supp Fig. 2A). Therefore, we performed a dose-response measurement of caspase activity using a caspase-cleavable fluorogenic probe, and increasing doses of radiation, in combination with increasing concentrations of IFN-β (Fig. 4A). The resulting heatmap revealed a dose-dependent increase in fluorescent signal derived from caspase 3/7 activity following irradiation. In contrast, IFN-β appeared to exert a protective effect, reducing the observable fluorescent signal upon treatment (Fig. 4A). This observation was further strengthened by quantifying apoptotic cells from IFN-β-treated organoids, based on Annexin V/Propidium Iodide (AnV+/PI+) positivity using flow cytometry and an identical gating strategy (Fig. 4B and C). As such, the cytokine was able to reduce the percentage of early apoptotic cells (AnV^+^/PI^−^), accompanied by an increase in live cells (double negative, AnV^−^/PI^−^). Still, no changes were observed in double positive (late apoptotic, or secondary necrotic cells), nor in singly stained PI^+^ cells (necrotic cells), suggesting the protective function of IFN-β might be exclusive to the apoptotic pathway.

**Figure 4.**
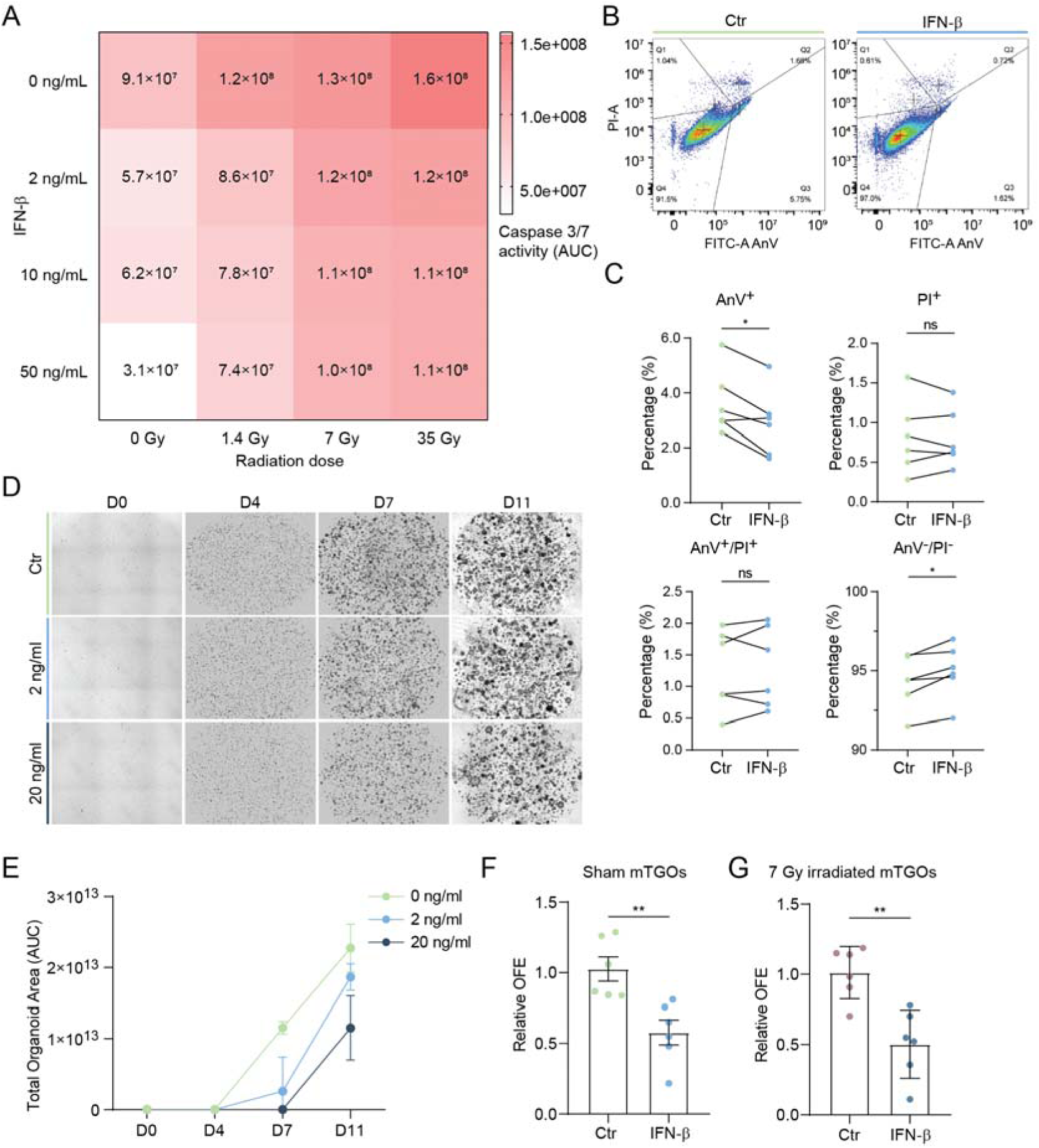
IFN-β exposure is protective for apoptotic cell death but impairs thyroid stem progenitor cell self-renewal. (A) Heatmap of caspase 3/7 activity in mTGOs following irradiation and IFN-β treatment with escalating doses. Data represents the area under the curve after 2 days (n = 6 animals/condition). (B) Representative dot plots, and (C) FACS analysis of AnV-FITC/PI staining in 11-day-old mTGO-derived cells following IFN-β treatment from D5 (right) and vehicle treated (left, Ctr). Data shown are percentage (%) of positive cells. Each dot represents a different biological replicate. Lines connect the same biological replicate across conditions. (D) Representative images of mTGOs cultured from D0 with 2 or 20 ng/ml IFN-β (and controls) at D0, D4, D7 and D11. (E) Quantification of the organoid area of mTGOs cultured from D0 with 2 or 20 ng/ml IFN-β (and controls) measured at D0, D4, D7 and D11. (F) Quantification of OFE in sham-irradiated mTGOs and (G) 7 Gy-irradiated mTGOs after IFN-β treatment from D5 to D11. Data are shown as mean ± SEM. Group differences were evaluated with Student *t-*test. n=3-6 animals/condition. \**p* < 0.05, \*\**p* < 0.01

Next, to characterize the impact of IFN-β on thyroid cell growth, the cytokine was administered to isolated thyrocytes immediately following their seeding as single cells in Matrigel™ domes, at both low (2 ng/mL) and high (20 ng/mL) concentrations of IFN-β. Cultures were monitored using the IncuCyte live-cell imaging system at multiple time points (D0, D4, D7, and D11; Fig. 4D), revealing a dose-dependent effect of IFN-β, on organoid size, particularly evident at D7 and D11, but not at the earlier timepoints (e.g. D0 and D4 (Fig. 4E), nor when IFN-β was administered in more mature organoids treated from day 5 onwards for a shorter period of time (up to 48 hours, Supp. Fig. 2B-C, and Supplementary Fig. 3 A-B). In this setting, treatment in already formed organoids from D5 onwards only significantly negatively impacted organoid growth at the highest dose of 20 ng/mL after 6 days of treatment (Supplementary Fig. 3A and B).

Given the apparent long-term effect of IFN-β on thyroid cell self-renewal, we next investigated the persistent effect of IFN-β on thyroid stem cell capacity by treating mature organoids (5-day-old) with 2 ng/mL IFN-β, followed by passaging on day 11, but now in the absence of IFN-β, to measure any lasting effect on self-renewal ability (OFE). Seven days post-passaging, OFE was significantly reduced in IFN-β–exposed cultures relative to controls, indicating that despite its potential anti-apoptotic effects, IFN-β may negatively regulate stemness (Fig. 4D), suggesting potentially different effects on different cell types. Furthermore, the combination of radiation and IFN-β led to an even greater reduction in OFE, suggesting an additive detrimental effect on the regenerative capacity of thyroid organoids (Fig. 4E). Therefore, we hypothesize that IFN-β may help maintain thyroid cell survival following acute damage (e.g. following irradiation) but may come at the expense of thyroid stem progenitor cell self-renewal capacity.

## Discussion

Correct thyroid function and hormone production is essential to maintain body homeostasis, as these target almost all tissues in the body, influencing growth and metabolism [29]. Radiation-induced hypothyroidism is a well-documented yet mechanistically understudied complication in patients receiving radiotherapy for HNC [3], [6], [22], [30–32]. In this study, tissue-derived mTGOs have been employed to model and dissect the cellular and molecular events occurring in the thyroid following irradiation. We demonstrate that radiation impairs thyroid organoid formation and maintenance in a dose-dependent manner, suggestive of functional damage. Crucially, we recognized IFN-β-mediated signaling as a response central to radiation exposure, which, although partially protective against apoptosis, ultimately compromises stem cell self-renewal capacity and post-irradiation regenerative potential.

The possibility of using organoids to investigate these biological processes has been extensively demonstrated using Salivary Gland Organoids [24], [28]. Irradiation of mature mTGOs (5-day-old), as opposed to single cells, better mimics the complexity of the tissue and allows the tracking of both early and delayed responses, which are critical to understanding thyroid dysfunction post-radiotherapy. In the case of the thyroid, the observed decline in OFE following irradiation underscores the vulnerability of thyroid progenitor or stem-like cells to ionizing radiation. This effect was particularly pronounced at higher doses (7 and 15 Gy), with a complete ablation of organoid formation at 15 Gy. These findings are in line with earlier preclinical models of radiation-induced thyroid dysfunction [6], as well as clinical reports of the high radiosensitivity of thyroid tissue [3], [6], [22], and a decreased risk of secondary thyroid cancer formation following 20 Gy of irradiation [33].

Bulk RNAseq revealed time-dependent transcriptional changes, with substantial enrichment of inflammatory and IFN-I-related pathways after irradiation, suggesting the major form of cell death occurring in thyroid cells following irradiation to be caspase-dependent apoptosis *via* the intrinsic activation pathway. In addition, IFN-β signaling was markedly upregulated and sustained over time. These findings were reinforced by the elevated expression of ISGs, including MX1 and RIG-I, and the activation of the STAT1/pSTAT1 axis in irradiated mTGOs. Our data suggest that, while this response may initially serve a cytoprotective role, prolonged activation of IFN-β signaling may impair thyroid stem progenitor cell self-renewal and thereby their regenerative capacity, with significant consequences for thyroid tissue integrity.

IFN-β has been recognized as a leader cytokine for the inflammatory response that follows radiation in healthy tissues, potentially enhancing radiation-induced tumor clearance [12]. However, it has also been implicated as a candidate cytokine responsible for tissue damage that is later conducive to long-term side effects of radiotherapy in patients [28]. The functional dichotomy of IFN-β observed in this study is of translational relevance. On one hand, exogenous IFN-β treatment mitigated radiation-induced apoptosis, as evidenced by reduced caspase activity, suggesting a context-dependent protective effect. On the other hand, both low and high physiological concentrations of IFN-β significantly diminished organoid growth and stemness, particularly when administered at earlier stages during critical phases of thyroid organoid development.

Furthermore, the combination of IFN-β and radiation exposure exerted an additive negative effect on OFE, implying that IFN-β does not merely reflect a consequence of radiation damage but may actively participate in the deterioration of thyroid regenerative function. Clinically, our findings mirror observations of IFN-I-associated thyroid dysfunction in patients treated with IFN-α or IFN-β for viral infections or cancers, where autoimmune thyroiditis and hypothyroidism are prevalent outcomes [13–16]. Although we did not explore autoimmune markers in this model, the sustained inflammatory response observed may lay the groundwork for chronic immune-mediated thyroid injury *in vivo,* bridging the gap between the reported cases of radiotherapy-induced thyroid dysfunction and the toxic effect of IFN-I on this sensitive tissue. Our study adds mechanistic insight into the poorly understood link between radiotherapy and thyroid dysfunction. While DNA damage and reactive oxygen species generation are immediate consequences of radiation, our data highlight the importance of delayed responses, including inflammatory signaling and regeneration, in shaping long-term tissue outcomes. The identification of IFN-β as a mediator of reduced organoid viability suggests a double-edged role: potentially beneficial in oncologic contexts but deleterious for adjacent healthy thyroid tissue.

Supporting this dual role are recent works suggesting IFN-I signaling following irradiation enhances stem/progenitor cell activity in the salivary gland [28], and the intestine [34], suggesting organ or context-specific roles in normal tissue regeneration. Indeed, specific differences between stem cell populations, which are yet to be identified in the thyroid gland [21], may underlie these opposing effects.

There are several limitations to our study. First, while mTGOs are an advanced *ex vivo* model, they do not fully capture systemic immune responses or vascular interactions present *in vivo*. Second, the murine origin of the organoids may not fully replicate human thyroid biology or its response to IFN-β signaling. Future studies using human-derived thyroid organoids are essential to validate these findings. Additionally, the mechanistic dissection of the crosstalk between IFN-β signaling, apoptosis, and stem cell exhaustion is warranted to identify actionable targets for radioprotection and to explore potential protective interventions.

In conclusion, we demonstrate that radiation-induced IFN-β signaling plays a critical role in impairing thyroid gland regeneration by reducing cell viability and self-renewal capacity in organoid models. These findings offer a plausible molecular mechanism at least partially underlying the clinical observation of hypothyroidism following HNC radiotherapy and support the need for strategies aimed at preserving thyroid function in irradiated patients.

### Data availability

Data have been deposited in GEO.

## Acknowledgments

This work was supported by the Dutch Cancer Society (KWF) Grant Nr 12092.

## CRediT authorship contribution statement

**Rufina Maturi:** Conceptualization, Methodology, Investigation, Visualization, Writing – original draft, Project administration **Davide Cinat:** Conceptualization, Methodology, Visualization, Writing – original draft, Project administration. **Anne L. Jellema-de Bruin:** Investigation. **Gabriella De Vita**: Supervision, Funding acquisition. **Schelto Kruijff:** Supervision, Funding acquisition. **Rob P. Coppes:** Supervision, Writing – review & editing, Funding acquisition. **Abel Soto-Gamez:** Conceptualization, Investigation, Methodology, Visualization, Supervision, Writing – original draft, Writing – review & editing, Project administration

## Notes

https://www.ncbi.nlm.nih.gov/geo/query/acc.cgi?acc=GSE303892

### Competing Interest Statement

The authors have declared no competing interest.

## References

[1] M. D. Mody, J. W. Rocco, S. S. Yom, R. I. Haddad, and N. F. Saba, “Head and neck cancer,” The Lancet, vol. 398, no. 10318, pp. 2289–2299, Dec. 2021, doi: 10.1016/S0140-6736(21)01550-6.

[2] L. Barazzuol, R. P. Coppes, and P. van Luijk, “Prevention and treatment of radiotherapy-induced side effects,” Mol Oncol, vol. 14, no. 7, pp. 1538–1554, Jul. 2020, doi: 10.1002/1878-0261.12750.

[3] M. K. Rooney et al., “Hypothyroidism following Radiotherapy for Head and Neck Cancer: A Systematic Review of the Literature and Opportunities to Improve the Therapeutic Ratio.,” Cancers (Basel), vol. 15, no. 17, Aug. 2023, doi: 10.3390/cancers15174321.

[4] C. Contreras-Jurado, “Thyroid Hormones and Co-workers: An Overview,” 2025, pp. 3–16. doi: 10.1007/978-1-0716-4252-8_1.

[5] U. Beling and J. Einhorn, “Incidence of Hypothyroidism and Recurrences Following I131 Treatment of Hyperthyroidism,” Acta radiol, vol. 56, no. 4, pp. 275–288, Oct. 1961, doi: 10.3109/00016926109172822.

[6] B. A. Jereczek-Fossa, D. Alterio, J. Jassem, B. Gibelli, N. Tradati, and R. Orecchia, “Radiotherapy-induced thyroid disorders,” Cancer Treat Rev, vol. 30, no. 4, pp. 369–384, Jun. 2004, doi: 10.1016/j.ctrv.2003.12.003.

[7] S. A. Roman, “Endocrine tumors: evaluation of the thyroid nodule.,” Curr Opin Oncol, vol. 15, no. 1, pp. 66–70, Jan. 2003, doi: 10.1097/00001622-200301000-00010.

[8] D. Cinat, R. P. Coppes, and L. Barazzuol, “DNA Damage-Induced Inflammatory Microenvironment and Adult Stem Cell Response,” Front Cell Dev Biol, vol. 9, Oct. 2021, doi: 10.3389/fcell.2021.729136.

[9] D. Schaue, E. D. Micewicz, J. A. Ratikan, M. W. Xie, G. Cheng, and W. H. McBride, “Radiation and inflammation.,” Semin Radiat Oncol, vol. 25, no. 1, pp. 4–10, Jan. 2015, doi: 10.1016/j.semradonc.2014.07.007.

[10] M. McLaughlin et al., “Inflammatory microenvironment remodelling by tumour cells after radiotherapy,” Nat Rev Cancer, vol. 20, no. 4, pp. 203–217, Apr. 2020, doi: 10.1038/s41568-020-0246-1.

[11] R. R. Weichselbaum, H. Liang, L. Deng, and Y.-X. Fu, “Radiotherapy and immunotherapy: a beneficial liaison?,” Nat Rev Clin Oncol, vol. 14, no. 6, pp. 365–379, Jun. 2017, doi: 10.1038/nrclinonc.2016.211.

[12] E. Jöst, W. P. Roos, B. Kaina, and H. Schmidberger, “Response of pancreatic cancer cells treated with interferon-α or β and co-exposed to ionising radiation,” Int J Radiat Biol, vol. 86, no. 9, pp. 732–741, Sep. 2010, doi: 10.3109/09553002.2010.481321.

[13] D. T. W. Lui et al., “The Impact of Interferon Beta-1b Therapy on Thyroid Function and Autoimmunity Among COVID-19 Survivors,” Front Endocrinol (Lausanne), vol. 12, Sep. 2021, doi: 10.3389/fendo.2021.746602.

[14] F. Monzani, N. Caraccio, A. Dardano, and E. Ferrannini, “Thyroid autoimmunity and dysfunction associated with type I interferon therapy,” Clin Exp Med, vol. 3, no. 4, pp. 199–210, Apr. 2004, doi: 10.1007/s10238-004-0026-3.

[15] Y. Tomer, J. T. Blackard, and N. Akeno, “Interferon Alpha Treatment and Thyroid Dysfunction,” Endocrinol Metab Clin North Am, vol. 36, no. 4, pp. 1051–1066, Dec. 2007, doi: 10.1016/j.ecl.2007.07.001.

[16] Y. Oppenheim, Y. Ban, and Y. Tomer, “Interferon induced Autoimmune Thyroid Disease (AITD): a model for human autoimmunity,” Autoimmun Rev, vol. 3, no. 5, pp. 388–393, Jul. 2004, doi: 10.1016/j.autrev.2004.03.003.

[17] V. M. L. Ogundipe et al., “Generation and Differentiation of Adult Tissue-Derived Human Thyroid Organoids,” Stem Cell Reports, vol. 16, no. 4, pp. 913–925, Apr. 2021, doi: 10.1016/j.stemcr.2021.02.011.

[18] A. A. Kurmann et al., “Regeneration of Thyroid Function by Transplantation of Differentiated Pluripotent Stem Cells,” Cell Stem Cell, vol. 17, no. 5, pp. 527–542, Nov. 2015, doi: 10.1016/j.stem.2015.09.004.

[19] F. Antonica et al., “Generation of functional thyroid from embryonic stem cells,” Nature, vol. 491, no. 7422, pp. 66–71, Nov. 2012, doi: 10.1038/nature11525.

[20] M. Romitti et al., “Transplantable human thyroid organoids generated from embryonic stem cells to rescue hypothyroidism,” Nat Commun, vol. 13, no. 1, p. 7057, Nov. 2022, doi: 10.1038/s41467-022-34776-7.

[21] J. van der Vaart et al., “Adult mouse and human organoids derived from thyroid follicular cells and modeling of Graves’ hyperthyroidism,” Proceedings of the National Academy of Sciences, vol. 118, no. 51, Dec. 2021, doi: 10.1073/pnas.2117017118.

[22] J. W. Foley et al., “Gene expression profiling of single cells from archival tissue with laser-capture microdissection and Smart-3SEQ,” Genome Res, vol. 29, no. 11, pp. 1816–1825, Nov. 2019, doi: 10.1101/gr.234807.118.

[23] A. Soto-Gamez et al., “Mesenchymal stem cell-derived HGF attenuates radiation-induced senescence in salivary glands via compensatory proliferation,” Radiotherapy and Oncology, vol. 190, p. 109984, Jan. 2024, doi: 10.1016/j.radonc.2023.109984.

[24] D. Alterio et al., “Thyroid disorders in patients treated with radiotherapy for head-and-neck cancer: A retrospective analysis of seventy-three patients,” International Journal of Radiation Oncology*Biology*Physics, vol. 67, no. 1, pp. 144–150, Jan. 2007, doi: 10.1016/j.ijrobp.2006.08.051.

[25] V. M. L. Ogundipe, J. T. M. Plukker, T. P. Links, and R. P. Coppes, “Thyroid Gland Organoids: Current Models and Insights for Application in Tissue Engineering,” Tissue Eng Part A, vol. 28, no. 11–12, pp. 500–510, Jun. 2022, doi: 10.1089/ten.tea.2021.0221.

[26] D. Cinat, A. L. De Souza, A. Soto-Gamez, A. L. Jellema-de Bruin, R. P. Coppes, and L. Barazzuol, “Mitophagy induction improves salivary gland stem/progenitor cell function by reducing senescence after irradiation,” Radiotherapy and Oncology, vol. 190, p. 110028, Jan. 2024, doi: 10.1016/j.radonc.2023.110028.

[27] X. Peng et al., “Cellular senescence contributes to radiation-induced hyposalivation by affecting the stem/progenitor cell niche,” Cell Death Dis, vol. 11, no. 10, p. 854, Oct. 2020, doi: 10.1038/s41419-020-03074-9.

[28] J. F. Bromberg, C. M. Horvath, Z. Wen, R. D. Schreiber, and J. E. Darnell, “Transcriptionally active Stat1 is required for the antiproliferative effects of both interferon alpha and interferon gamma.,” Proceedings of the National Academy of Sciences, vol. 93, no. 15, pp. 7673–7678, Jul. 1996, doi: 10.1073/pnas.93.15.7673.

[29] D. Cinat et al., “Transposable Element Derepression-mediated IFN-I Signaling Enhances Stem/Progenitor Cell Activity After Irradiation,” Feb. 15, 2024. doi: 10.1101/2024.02.14.580306.

[30] R. Mullur, Y.-Y. Liu, and G. A. Brent, “Thyroid Hormone Regulation of Metabolism,” Physiol Rev, vol. 94, no. 2, pp. 355–382, Apr. 2014, doi: 10.1152/physrev.00030.2013.

[31] S. L. Turner, K. W. Tiver, and S. C. Boyages, “Thyroid dysfunction following radiotherapy for head and neck cancer,” International Journal of Radiation Oncology*Biology*Physics, vol. 31, no. 2, pp. 279–283, Jan. 1995, doi: 10.1016/0360-3016(93)E0112-J.

[32] L. Bernát and D. Hrušák, “Hypothyroidism after radiotherapy of head and neck cancer,” Journal of Cranio-Maxillofacial Surgery, vol. 42, no. 4, pp. 356–361, Jun. 2014, doi: 10.1016/j.jcms.2013.09.009.

[33] A. S. Randhawa, H. P. Yadav, R. P. S. Banipal, G. Goyal, P. Garg, and S. Marcus, “Functional and biochemical changes in the thyroid gland following exposure to therapeutic doses of external beam radiotherapy in the head-and-neck cancer patients,” J Cancer Res Ther, vol. 17, no. 4, pp. 1025–1030, Jul. 2021, doi: 10.4103/jcrt.JCRT_148_19.

[34] A. Berrington de Gonzalez et al., “Second Solid Cancers After Radiation Therapy: A Systematic Review of the Epidemiologic Studies of the Radiation Dose-Response Relationship,” International Journal of Radiation Oncology*Biology*Physics, vol. 86, no. 2, pp. 224–233, Jun. 2013, doi: 10.1016/j.ijrobp.2012.09.001.

[35] T. L. Lim et al., “Early Inflammation and Interferon Signaling Direct Enhanced Intestinal Crypt Regeneration after Proton FLASH Radiotherapy,” Aug. 19, 2024. doi: 10.1101/2024.08.16.608284.

