## Supplementary Table 1 for "Radiation-induced interferon-I response impairs thyroid organoid function"

| **Gene** | **Forward Primer 5’-3’** | **Reverse Primer 5’-3’** |
| --- | --- | --- |
| Ywhaz | TTACTTGGCCGAGGTTGCT | TGCTGTGACTGGTCCACAAT |
| Oas2 | TTGAAGAGGAATACATGCGGAA | GGGTCTGCATTACTGGCACTT |
| Ifit1 | CTGAGATGTCACTTCACATGGAA | GTGCATCCCCAATGGGTTCT |
| Cxcl10 | GCCGTCATTTTCTGCCTCA | CGTCCTTGCGAGAGGGATC |
| Ccl5 | TGCCCACGTCAAGGAGTATTTC | AACCCACTTCTTCTCTGGGTTG |
| Ddx58 (Rig-I) | AAGAGCCAGAGTGTCAGAATCT | AGCTCCAGTTGGTAATTTCTTGG |

**Supplementary Table 1.** qPCR primer sequences.
