## Supplementary Fig.1 for "Radiation-induced interferon-I response impairs thyroid organoid function"

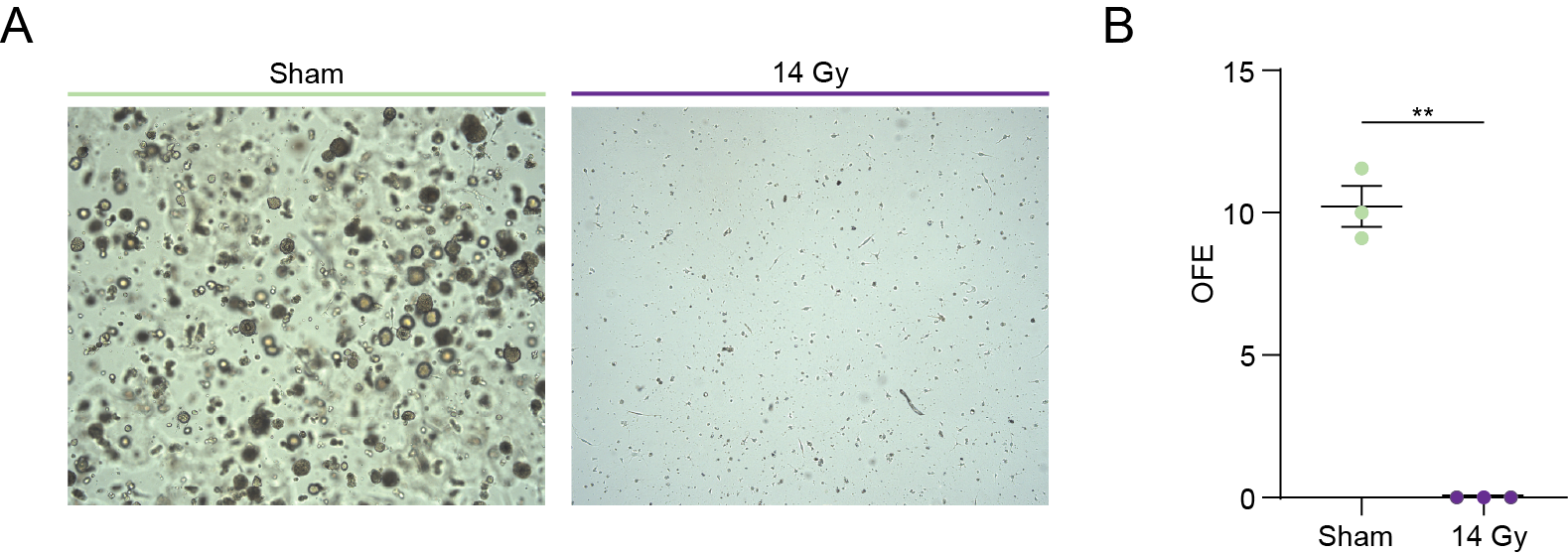

Sham	15 Gy

Sham

15 Gy

**Supplementary Figure 1. *In vivo irradiation abolishes thyroid cell ability to grow organoids.*** (A) Representative images of 7-day-old mTGOs obtained from 15 Gy irradiated mice (right) and controls (left). (B) Organoid quantification shown as absolute OFE. Data are shown as mean ± SEM. N=3. ***p* < 0.01
