## Supplementary Fig.2 for "Radiation-induced interferon-I response impairs thyroid organoid function"

**A	B	C**

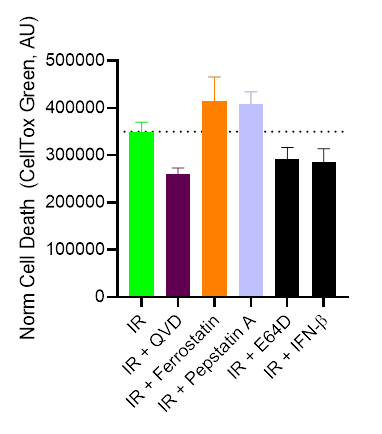

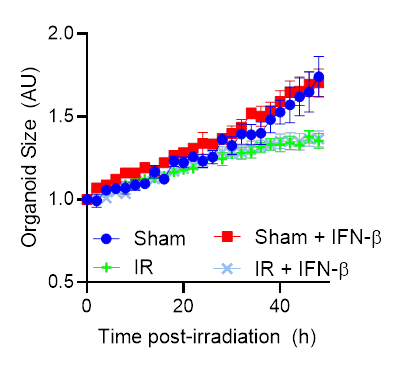

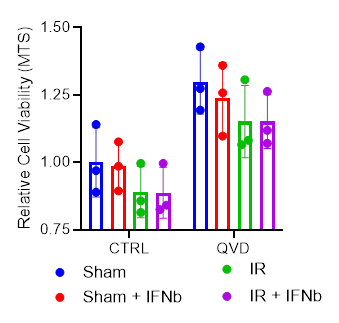

**Supplementary Figure 2. Effect of irradiation and short-term exposure (48h) to IFN-β in thyroid gland organoids.** Thyroid gland organoid cultures (D5) were irradiated (7Gy) and treated with to IFN*-β* for 48 hours. (A) The effect of irradiation and short-term exposure to IFN-β on cell death was tested in the presence of known cell death inhibitors (QVD: caspase-dependent apoptosis inhibitor, Ferrostatin: ferroptosis inhibitor, Pepstatin A: cathepsins D and E-induced death inhibitor, and E64D: cathepsin B and L-, and calpains-induced death, all tested at 10 µM). The early effects of irradiation and IFN-β (up to 48h) were measured on (B) organoid size and (C) cell viability. Data are shown as mean ± SEM. N=3. ***p* < 0.01.
