## Supplementary Fig.3 for "Radiation-induced interferon-I response impairs thyroid organoid function"

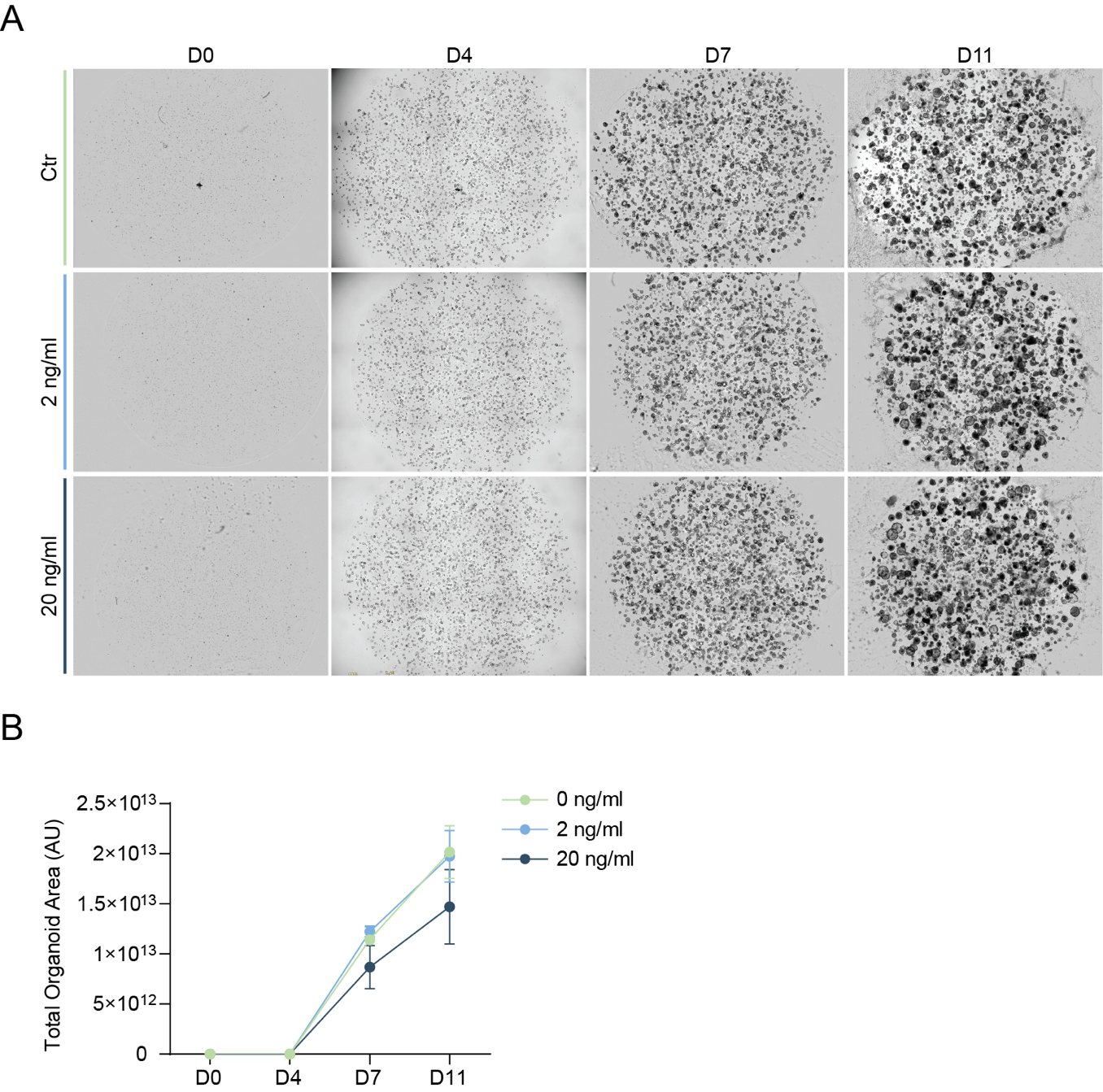

**Supplementary Figure 3. Long-term *IFN-β exposure reduces thyroid cell self-renewal ability.*** (A) Representative images of mTGOs cultured from D5 with 2 or 20 ng/ml IFN-β (and controls) at D0, D4, D7, D11, using the Organoid module. (B) Quantification of the organoid area calculated at D0, D4, D7, D11, with IncuCyte® S3, of mTGOs cultured from D5 with 2 or 20 ng/ml IFN-β (and controls). Data are shown as mean ± SEM. N=3
